# Mechanism of MCUB-dependent inhibition of mitochondrial calcium uptake

**DOI:** 10.1101/2024.11.25.625228

**Authors:** Neeraj K. Rai, Anthony M. Balynas, Melissa JS MacEwen, Ashley R. Bratt, Yasemin Sancak, Dipayan Chaudhuri

**Author notes:** **Corresponding author:** Dipayan Chaudhuri, MD, PhD, 95 South 2000 East, Building 500 (CVRTI), Salt Lake City, UT 84112.

## Abstract

Mitochondrial Ca^2+^ levels are regulated to balance stimulating respiration against the harm of Ca^2+^ overload. Contributing to this balance, the main channel transporting Ca^2+^ into the matrix, the mitochondrial Ca^2+^ uniporter, can incorporate a dominant-negative subunit (MCUB). MCUB is homologous to the pore-forming subunit MCU, but when present in the pore-lining tetramer, inhibits Ca^2+^ transport. Here, using cell lines deleted of both MCU and MCUB, we identify three factors that contribute to MCUB-dependent inhibition. First, MCUB protein requires MCU to express. The effect is mediated via the N-terminal domain (NTD) of MCUB. Replacement of the MCUB NTD with the MCU NTD recovers autonomous expression but fails to rescue Ca^2+^ uptake. Surprisingly, mutations to MCUB that affect interactions with accessory subunits or the conduction pore all failed to rescue Ca^2+^ uptake, suggesting the mechanism of inhibition may involve global rearrangements. Second, using concatemeric tetramers with varying MCU:MCUB ratios, we find that MCUB incorporation does not abolish conduction, but rather inhibits Ca^2+^ influx proportional to the amount of MCUB present in the channel. Reducing rather than abolishing Ca^2+^ transport is consistent with MCUB retaining the highly-conserved selectivity filter DIME sequence. Finally, we apply live-cell Förster resonance energy transfer to establish that the endogenous stoichiometry is 2:2 MCU:MCUB. Taken together, our results suggest MCUB preferentially incorporates into nascent uniporters, and the amount of MCUB protein present linearly correlates with the degree of inhibition of Ca^2+^ transport, creating a precise, tunable mechanism for cells to regulate mitochondrial Ca^2+^ uptake.

## INTRODUCTION

Ca^2+^ entry into the mitochondrial matrix helps control fuel preference for respiration, stimulates the citric acid cycle, and helps regulate oxidative phosphorylation to maintain cellular bioenergetics (Lee et al., 2023). However, various forms of injury and stress induce Ca^2+^ overload, a pathological process that can ultimately lead to cell death. Thus, cellular control of mitochondrial Ca^2+^ fluxes is essential to balance bioenergetic needs against preventing mitochondrial dysfunction.

To enter the matrix, Ca^2+^ activates and travels through a channel known as the mitochondrial Ca^2+^ uniporter, a multiprotein complex that resides in the inner mitochondrial membrane (IMM) (Balderas et al., 2023; Kamer & Mootha, 2015; MacEwen & Sancak, 2023). This complex contains a tetramer of pore-forming subunits (MCU), a gatekeeping dimer composed of MICU1 and either MICU2 or MICU3, and an accessory subunit (EMRE) that encages the pore, holding it open and bridging interactions with the MICU proteins. Interactions with additional proteins, such as MCUR1, can regulate channel behavior.

Intriguingly, Raffaello et. al. described a protein, MCUB, that shares ∼50% amino acid similarity to MCU, but appears to inhibit Ca^2+^ uptake in a dominant negative manner when it replaces one or more MCU subunits in the pore (Raffaello et al., 2013). MCUB is substantially upregulated during stress in a manner that suggests that its physiological function is to reduce the risk of toxic Ca^2+^ overload during pathological processes (Huo et al., 2020; Lambert et al., 2019).

The mechanism by which MCUB inhibits Ca^2+^ uptake remains unresolved. Purified MCUB does not form Ca^2+^ channels in isolation in planar lipid bilayers, and mutations to MCU that mimic MCUB abolish Ca^2+^ uptake, implying that MCUB may entirely block ion transport (Raffaello et al., 2013). However, there is also evidence suggesting MCUB-containing channels can transport Ca^2+^. MCUB possesses the highly-conserved DIME motif that binds Ca^2+^ at the channel mouth, and purified MCUB in bilayers does conduct Na^+^. In this light, further studies have suggested other mechanisms for inhibition, such as alterations in channel electrostatics, pore structure, or interactions with accessory subunits (Colussi & Stathopulos, 2023).

How MCUB assembles into the channel, and whether it does so with fixed or variable stoichiometry have also been critical open questions. MCUB mimics MCU in its domain structure, encoding an N-terminal domain (NTD) that resembles a β-grasp fold, a coiled-coil domain (CCD), and two transmembrane domains (TMD). Recent studies have shown that the isolated MCUB NTD (NTb) binds preferentially with the MCU NTD (NTa), but it remains unclear how this contributes to channel architecture (S. K. Lee et al., 2016; Noble et al., 2024).

Here, we tackle these open questions taking advantage of a HEK-293T cell line in which both MCU and MCUB have been deleted by gene editing. First, we find that MCUB will not express in the absence of MCU. Second, increasing the amount of MCUB relative to MCU in each channel tetramer leads to a graded decrease in Ca^2+^ uptake, suggesting MCUB does not fully abolish Ca^2+^ uptake. Finally, we find that the native channel stoichiometry is 2:2 MCU:MCUB, with assembly driven by NTa-NTb interactions. These results suggest a simple assembly mechanism that allows precise, linear regulation of mitochondrial Ca^2+^ uptake.

## METHODS

### Plasmids

Plasmids for expression in HEK-293T cells were made by Gibson assembly using NEBuilder HiFi DNA Assembly Kit and gBlocks (Integrated DNA Technologies) or human codon-optimized constructs were synthesized by Twist Biosciences. All constructs were verified by sequencing. MCU-mCerulean and MCU-Flag constructs have been described previously (Balderas et al., 2022; Chaudhuri et al., 2013).

### Cell culture and generation of stable lines

MCU/MCUB double knockout (DKO) HEK-293T cells were a kind gift from the laboratory of Vamsi Mootha (Sancak et al., 2013). HEK-293T cells were cultured in Dulbecco’s Modified Eagle Medium (DMEM) (ThermoFisher), supplemented with 10% fetal bovine serum and the antibiotics penicillin (100 I.U./ml) and streptomycin (100 μg/ml), at 37°C and 5% CO2. Stable cell lines were generated by lentiviral transduction followed by selection with 1.5 µg/mL puromycin.

### Western Blotting

Total protein was extracted from cells with radioimmunoprecipitation assay (RIPA) lysis buffer supplemented with protease and phosphatase inhibitors (Thermo) as described previously (Sommakia et al., 2017). BCA assay was used for the measurement of protein concentration. Then, 20-40 µg of protein was run on 4-12% Bolt SDS-PAGE gels (Thermo) and transferred onto PVDF membrane for blocking and immunostaining. We used the following antibodies: EMRE (A300-BL19208, Bethyl, 1:1000), FLAG HRP (A8592, Sigma, 1:1000), GAPDH (2118S, CST, 1:1000), HA (3724S, CST, 1:100).

### Mitochondrial Ca^2+^ uptake

We measured Ca^2+^ uptake using fluorescent sensors as previously described (Chaudhuri et al., 2016; Eberhardt et al., 2022). Briefly, cells were harvested and resuspended in a medium containing 125□mM KCl, 2□mM K_2_HPO_4_, 1□mM MgCl_2_, 20□mM HEPES, 5□mM succinate, 0.01% digitonin, and 1□μM Oregon Green BAPTA 6F (OGB6F) at a concentration of 10^7^ cells/mL Imaging was performed in clear-bottom 96-well plates on a Cytation 5 microplate reader. 10^6^ cells were aliquoted in a well and OGB6F fluorescence was sampled at 490 nm excitation and 515 nm emission at room temperature, before and after a 25 µM Ca^2+^ bolus injection.

### Förster resonance energy transfer (FRET)

Flow cytometric FRET measurements were performed as described previously (Balderas et al., 2022; S. R. Lee et al., 2016). The use of the donor (E_D_) and acceptor (E_A_) FRET efficiency ratios (E_A_/E_D_) to assess stoichiometry follows the analysis described previously (Ben-Johny et al., 2016). Cells were imaged 1-3 days after transfecting MCU/MCUB DKO or HEK-293T cells in 6-well plates. Prior to resuspension and analysis, cells were incubated with 100 µM cycloheximide for ∼1-2 hours to prevent inaccurate measurements due to incomplete protein synthesis of fluorophores. Flow cytometry was done on a BD FACS Canto Analyzer, where signals were collected for the following: (i) 408 nm laser excitation and 450/50 nm emission for mCerulean; (ii) 408 nm excitation and 525/50 nm emission for FRET; and (iii) 488 nm excitation and 530/30 nm emission for mVenus. Single live-cell populations were gated based on forward and side scatter parameters. Signals from untransfected cells were used for background correction. Background-corrected signals are denoted as S_Ven_, S_Cer_, and S_FRET_ for the excitation/emission pairs 488/530, 408/450, and 408/525 nm, respectively. Cells transfected with mt-mVenus alone were used to calculate the crosstalk ratio between S_Ven_ and S_FRET_; cells transfected with mt-mCerulean alone were used to calculate crosstalk ratios between S_Cer_ and S_FRET_ and between S_Cer_ and S_Ven_. These ratios were used to obtain the amount of fluorescence in the FRET channel due to Cerulean crosstalk (Cer_direct_), Venus crosstalk (Ven_direct_) and Venus FRET (Ven_FRET_). Signals from mitochondria-targeted Cerulean and Venus dimers with linkers of different lengths (see Fig. 5C) were used to obtain fluorophore-(f_ven_/f_cer_) and instrument-(g_cer_/g_ven_) specific constants. We then selected transfected cells with high Cerulean and Venus fluorescence to calculate the E_A_/E_D_ ratio using these parameters:

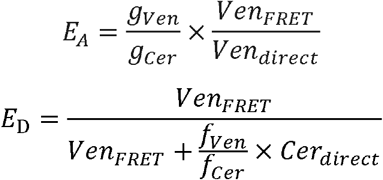

Data were analyzed using FlowJo and Microsoft Excel software.

## RESULTS

### MCUB protein expression requires the presence of MCU

To avoid confusion, we refer to the mitochondrial Ca^2+^ uniporter channel complex as the uniporter, and to its components by their gene/protein names (e.g. MCU, MCUB).

To investigate how MCUB inhibits Ca^2+^ uptake through the uniporter, we obtained HEK-293T cells from Dr. Vamsi Mootha in which MCUB and MCU had both been deleted (MCU/MCUB DKO) using TALEN technology, leading to frameshift indels and early termination (Sancak et al., 2013). Unfortunately, none of the current commercially-available MCUB antibodies produce specific bands (Huo et al., 2020), so we further validated the cell line by examining mRNA transcripts for nonsense-mediated decay. We found that MCU and MCUB transcript levels were 35±2% and 23±2% (n=3) of wild-type controls, respectively. Accessory uniporter subunits EMRE and MICU1 were only modestly affected (70±5% and 132±11% of wild-type, n=3)

Next, to assess the behavior of MCUB protein in the DKO cells, we tagged MCU and MCUB with Flag and HA tags, respectively. The recombinant MCUB-HA was transfected alone and co-expressed with MCU-Flag in DKO cells. The day following transfection, we collected lysates for immunoblotting. Notably, MCUB expressed when co-transfected with MCU but failed to express when transfected alone (Fig. 1), revealing that an interaction with MCU is necessary for robust MCUB expression. This is consistent with prior data from MCU KO cells, where MCUB expression was absent, and from hearts, where very low levels of MCU are present, leading to absent MCUB protein expression despite the presence of MCUB mRNA (Huo et al., 2020; Lambert et al., 2019; Luongo et al., 2015; Sancak et al., 2013). Taken together, these data suggest MCUB is unstable and cleared if MCU is not present via a post-translational mechanism.

**Figure 1.**
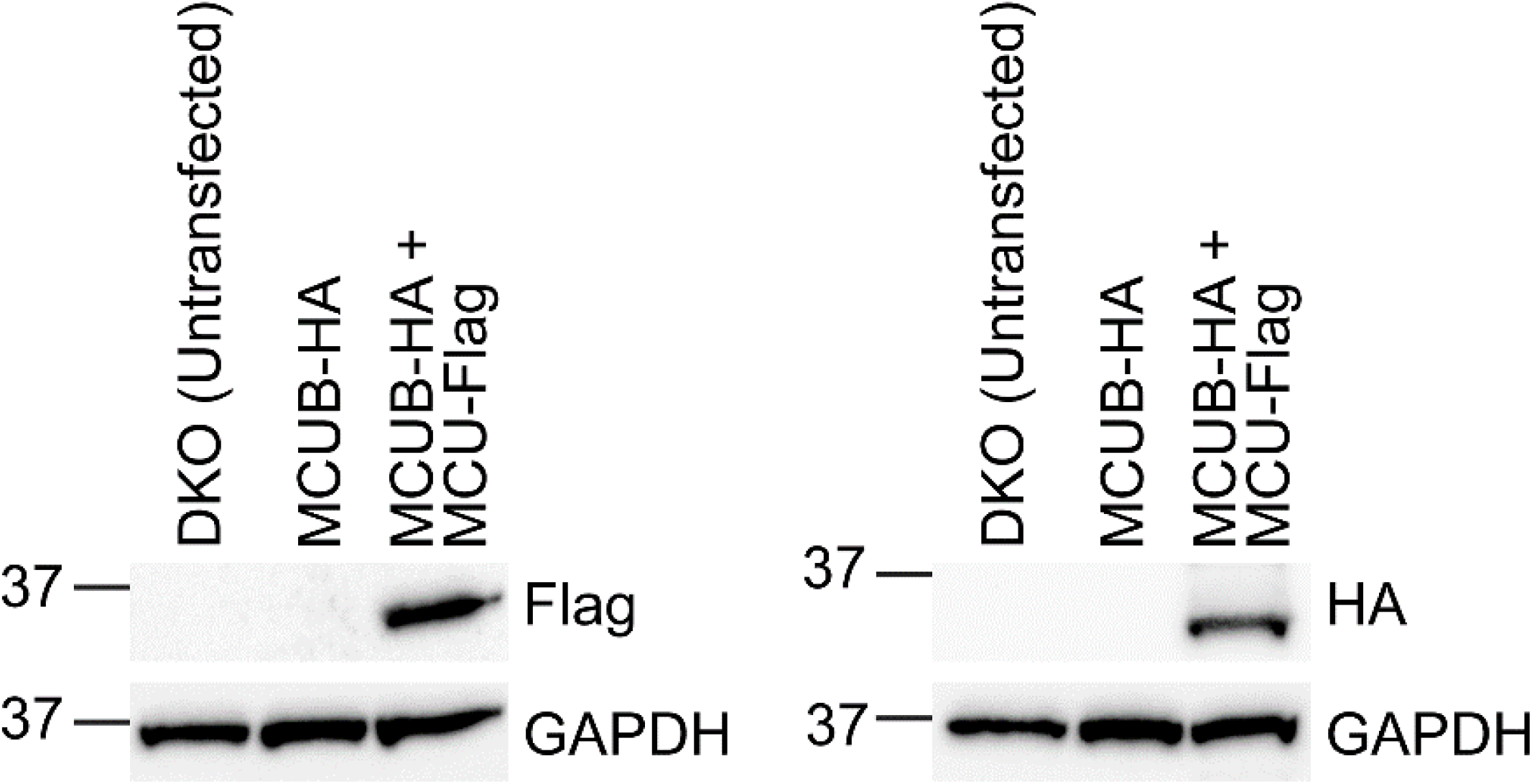
MCUB does not express without MCU. Western blot using MCU/MCUB DKO HEK 293T cells. Untransfected cells (DKO) or cells transfected with MCUB-HA alone or co-transfected with MCUB-HA and MCU-Flag were lysed and probed for HA, Flag, or GAPDH loading control. Identical samples were loaded in different gels because Flag/HA staining could not be entirely stripped away.

### MCU N-terminal domain helps MCUB to express in MCU-deficient cells

To examine why, despite sharing ∼50% sequence identity with MCU, MCUB does not express by itself, we pursued a mutagenesis strategy (Fig. 2). We introduced mutations in MCUB to mimic MCU, and examined whether these constructs would express in isolation in MCU/MCUB DKO cells. A V242E substitution in MCUB might impact the kinetics of Ca^2+^ permeation by increasing surface electrostatic potential, while the S229A substitution may alter hydrophobic interactions both with neighboring MCUB and EMRE (Colussi & Stathopulos, 2023; Raffaello et al., 2013). Since EMRE is essential for conduction of functional MCU channels, we also mutated the EMRE dependent domain (EDD) to mimic MCU more closely (MacEwen et al., 2020; Van Keuren et al., 2020). Finally, we replaced the soluble (matrix-resident) N-terminal domain of MCUB (NTb) with MCU N-terminal domain (NTa), since these domains are implicated in channel interactions (Balderas et al., 2022; Dong et al., 2017; S. K. Lee et al., 2016; Lee et al., 2015; Noble et al., 2024). We transfected each of these constructs into MCU/MCUB DKO cells, and assayed them by Western blot a day later. Intriguingly, only the MCUB mutant having the MCU NTD was expressed (MCUB-NTa, Fig. 2B). This suggests that full length MCUB fails to express unless MCU is present due to interactions between NTa-NTb and further implies that without binding to MCU, MCUB will not be expressed in clusters alone. Such a mechanism allows MCUB to maximize its inhibition by only forming hetero-oligomeric MCU-MCUB clusters.

**Figure 2.**
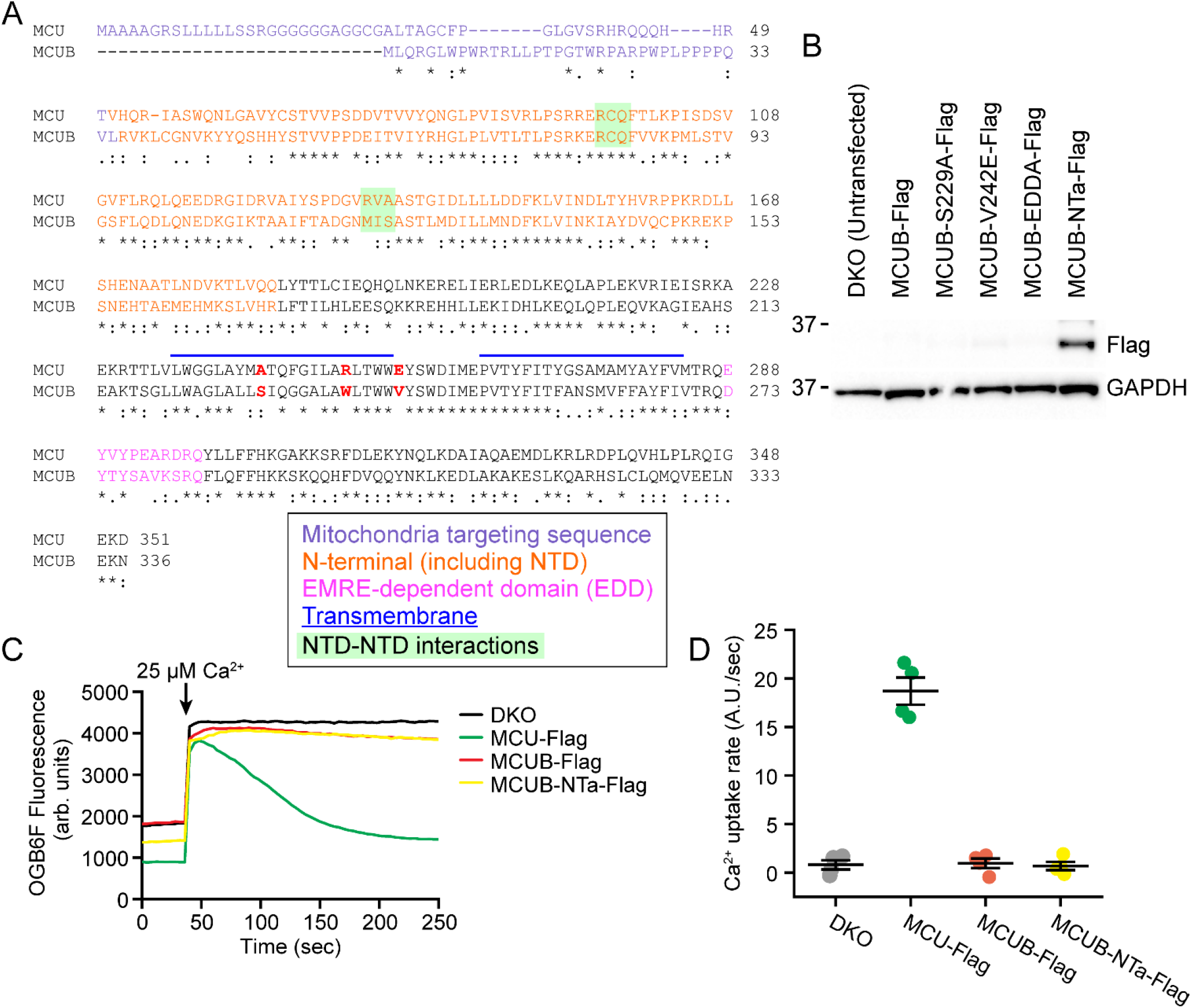
The MCUB NTD prevents expression of isolated MCUB. **A**. Alignment of human MCU and MCUB. Domains and point mutations (red) that were transplanted from MCU to MCUB to make heterologous constructs listed in 2B-D and Figure 3 are indicated. The MCU NTD (NTa) extends from amino acid ∼72-165, while the MCUB NTD is ∼58-151. **B**. Western blot analysis of Flag-tagged MCUB mutants. Only MCUB mutants containing NTa expressed in isolation when transfected into MCU/MCUB DKO cells. **C**. Exemplars, Ca^2+^ uptake assay. MCU/MCUB DKO cells were transfected with the indicated constructs and assayed the next day. Cells were permeabilized, and Oregon Green BAPTA 6F (OGB6F) fluorescence measured during a 25 µM Ca^2+^ pulse. MCUB-NTa mutants express but do not rescue Ca^2+^ uptake. **D**. Summary of uptake rates (n=4).

### A range of mutations to the MCUB transmembrane domains fails to confer Ca^2+^ uptake

Although purified MCUB was unable to form Ca^2+^ channels in lipid bilayers, this has not been directly tested in the intact mitochondria. The finding that MCUB-NTa can be expressed in isolation allowed us to investigate how MCUB prevents Ca^2+^ conduction. While mutating key residues in MCU to their corresponding MCUB cognates impaired uniporter Ca^2+^ conduction (Raffaello et al., 2013), we used MCUB NTa to test the opposite scenario: whether we could create a Ca^2+^ channel with only MCUB. Therefore, we next tested whether MCUB-NTa expressed alone would be capable of mitochondrial Ca^2+^ uptake. For this purpose, we use an extra-mitochondrial Ca^2+^ clearance assay, using the fluorescent membrane-impermeable Ca^2+^ indicator Oregon Green BAPTA 6F (OGB6F). In this assay, cells are treated with digitonin, which permeabilized most membranes but leaves intact the mitochondrial inner membrane. A 25 µM pulse of Ca^2+^ causes a rapid increase in fluorescence, which subsequently subsides as Ca^2+^ enters the mitochondria. Using this assay, MCU/MCUB DKO cells expressing MCU alone displayed robust Ca^2+^ uptake, whereas no uptake was visible in MCU/MCUB DKO cells (Fig. 2C-D). However, DKO cells expressing MCUB-NTa failed to transport Ca^2+^ into mitochondria. These results establish that the MCUB protein fails to take up Ca^2+^ even when robustly expressed, suggesting that alterations to the transmembrane domains prevent formation of a permeant channel.

To examine whether interactions with EMRE, selectivity filter, or other structural variants were responsible for the absence of conduction in MCUB, we systematically made a range of mutations to the MCUB-NTa transmembrane segments to bring these closer to the MCU counterparts. First, we tested them for expression by Western blot (Fig. 3A). We then measured the Ca^2+^ uptake of these MCUB mutants (Fig. 3B). Surprisingly, all the mutants failed to restore the Ca^2+^ uptake. This suggests that the inhibition of Ca^2+^ uptake is not specified by one or a few mutations near the selectivity filter, but by distributed changes that likely more subtly alter Ca^2+^ transport.

**Figure 3.**
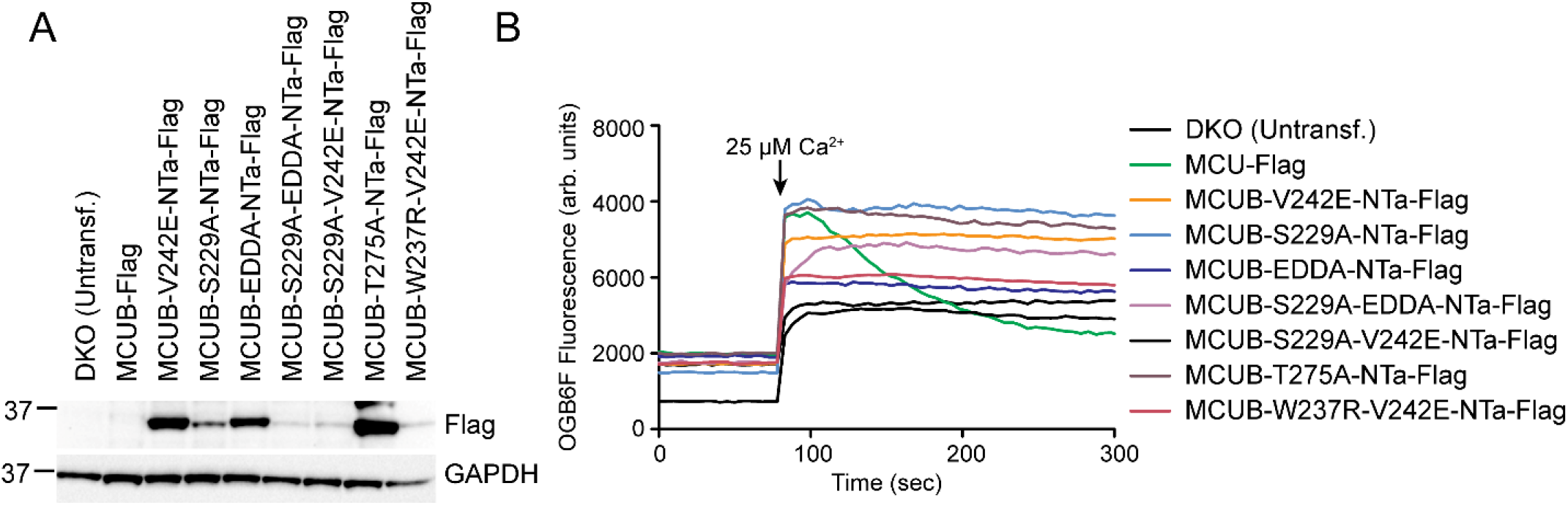
Various MCUB mutants fail to rescue Ca^2+^ uptake when expressed in isolation. **A**. Western blot showing expression of mutants as in Fig. 2B. Transplanted domains and point mutants as in Fig. 2A. Note that mutants with more extensive changes have reduced expression. **B**. Ca^2+^ uptake assay as in Fig. 2C. All the mutants failed to produce robust Ca^2+^ uptake.

### Increasing MCUB occupancy within uniporter channels progressively reduces but does not abolish Ca^2+^ uptake

A central open question is whether MCUB incorporation abolishes or slows Ca^2+^ flux through individual channels. Because each uniporter channel is a tetramer of MCU/MCUB subunits, assaying the contribution of MCUB within individual channels presents a technical challenge. We take an innovative approach inspired by the architecture of plasma membrane voltage-gated channels. Whereas voltage-gated K^+^ channels, like the uniporter, are formed by tetramers of single subunits, voltage-gated Na^+^ and Ca^2+^ channels have four such subunits tethered together by linkers into a single gene. We use a similar strategy to join different combinations of MCU and MCUB into single peptides (Fig. 4A-B). Prior attempts at such tetramer concatenation may have failed because short linkers were used, whereas the N- and C-termini of adjacent MCU subunits in the cryo-EM structures are ∼50 Å apart (Payne et al., 2020; Wang et al., 2019). To overcome this potential problem, we used a 28 amino acid highly-flexible linker (∼60-70 Å) derived from the TonB-dependent hemoglobin/transferrin/lactoferrin family receptor from *Serratia marcescens* (HasR) (Wojtowicz et al., 2016). We term the single protein composed of four identical linked MCU subunits MCUaaaa. MCUaaaa expresses as a band on Western blot 4x the monomer size (Fig. 4C). When expressed in MCU/MCUB DKO cells, MCUaaaa binds EMRE and produces uniporter-like Ca^2+^ uptake, confirming functionality (Fig. 4C-D)

**Figure 4.**
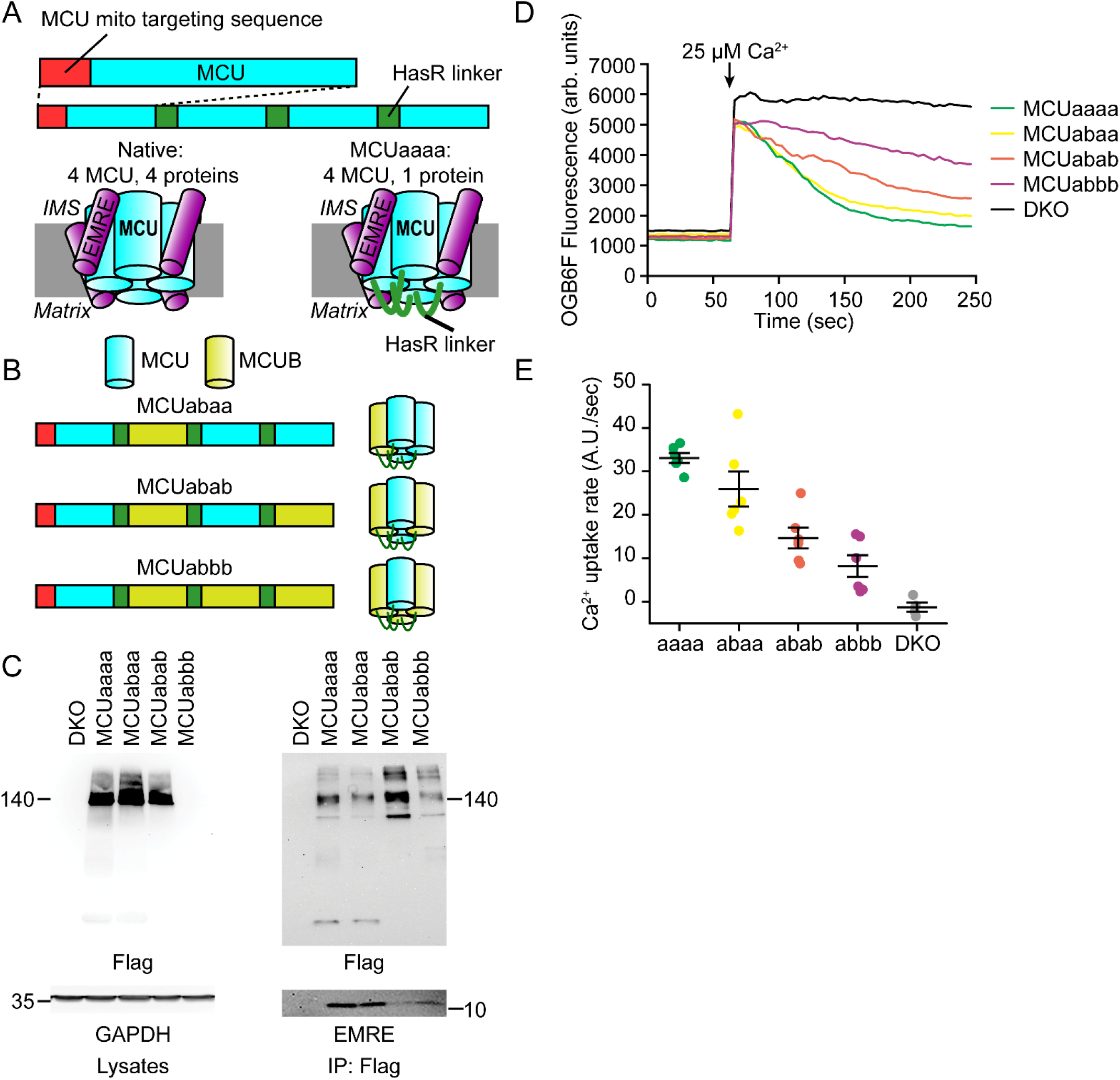
MCUB inhibits Ca^2+^ uptake in proportion to its channel occupancy. **A**. Concatenation strategy. We arrayed 3 mature (no targeting sequence) MCU sequences after a full-length MCU, connecting each repeat with a long HasR linker. Thus, the entire tetrameric pore is encoded by one gene. **B**. We created channels with different MCUB occupancy by substituting the MCUB sequence into the concatemers as indicated. **C**. MCU concatemers express and co-immunoprecipitate with EMRE. Left, lysates as in Fig. 2B of MCU/MCUB DKO cells with different concatemers. Right, lysates were immunoprecipitated with anti-Flag beads, and eluates stained for Flag and EMRE. MCUabbb expression is weak but present. Bands at lower molecular weights for MCUaaaa and MCUabaa may represent low levels of free monomer. **D, E**. Ca^2+^ uptake assay and summary as in Fig. 2C, D (n=4-6). There is a graded inhibition of Ca^2+^ uptake with greater MCUB occupancy within each tetramer.

Next, we generated constructs encoding tetrameric concatemers with different ratios of MCU to MCUB. To construct these concatemers, we utilize monomers of MCU and MCUB arranged in a head-to-tail configuration (MCUaaaa, MCUabaa, MCUabab, and MCUabbb), and subsequently established stable lines in MCU/MCUB DKO cells. The cells produced a fusion protein with a molecular weight consistent with a tetrameric uniporter (Fig. 4C, though MCUabbb expressed weakly), and all these concatemers interacted with EMRE (Fig. 4C). These results suggest that the concatemers assemble into functional uniporter tetramers. We next evaluated how MCUB stoichiometry altered conduction through the uniporter. We found that an increase in MCUB ratio within the tetramer produced a graded decrease in mitochondrial Ca^2+^ uptake (Fig. 4D-E). Though the response for the MCUabbb construct may have been affected by its poorer expression, nevertheless our data shows that increasing MCUB within a tetramer reduces global Ca^2+^ uptake.

### MCU assembles with MCUB in a two-to-two ratio

To determine the native stoichiometry of MCUB-containing uniporters, we turned to Förster resonance energy transfer (FRET). This spectroscopic phenomenon can assay interactions over short distances due to the sixth-power dependence of energy transfer on the distance between acceptor and donor fluorophores. Tagging candidate proteins with genetically-encoded fluorophores allows interactions between proteins to be assayed in live cells. For the mVenus-mCerulean pair used here, the Förster radius for 50% FRET efficiency is ∼5 nm (Bajar et al., 2016). Notably, under high-expression conditions, where all acceptor and donor fluorophores are within complexes and produce high signal-to-noise fluorescence, FRET analysis can elucidate the stoichiometry of these complexes (Ben-Johny et al., 2016). FRET causes opposite changes in fluorescence emission for the acceptor and donor molecule. Increasing FRET reduces donor fluorescence intensity, as more energy is transferred to the acceptor rather than photon emission. Conversely, increasing FRET increases acceptor fluorescence emission. These opposite changes can be quantified by measuring FRET efficiency from the standpoint of either donor or acceptor, with the donor-centric measure assaying the reduction in donor emission due to FRET, while the acceptor-centric measure quantifies the fractional increase in sensitized emission due to FRET. To reveal how this can be used to determine stoichiometry, we illustrate the process using uniporters with different MCU:MCUB composition (Fig. 5A). Here we tag MCU with mCerulean, the FRET donor, while MCUB is tagged with mVenus, the acceptor. When there is one MCU but 3 MCUBs, there is substantial FRET when measured from the perspective of the donor (E_D_), but much weaker FRET from the acceptor perspective (E_A_), since each MCUB receives only a fraction of the energy transfer. In this system, the ratio of donor molecules to acceptor molecules is calculated by the E_A_/E_D_ ratio (∼0.33). Conversely, for a channel with 3 MCU and 1 MCUB, E_D_ is low because there is only one acceptor for FRET, whereas E_A_ is high, because that acceptor is receiving energy from 3 different donors. In this case, the E_A_/E_D_ ratio will be 3. Finally, if equal numbers of MCU and MCUB are in each complex, then we expect E_A_ = E_D_ with a ratio of 1. However, this is the situation for an isolated channel, and the E_A_/E_D_ measured for each cell will report an average for all uniporters within that cell (Fig. 5B). To examine the distribution in live cells, we need to ensure high expression of both molecules for robust E_A_/E_D_ quantification (green shaded region in Fig. 5C-D). Under this condition, measurement of the E_A_/E_D_ ratio in live cells will produce one of two outcomes. If uniporters assemble with variable MCU:MCUB composition, then the E_A_/E_D_ ratio from each cell will be a weighted average of complexes in that cell with 3:1, 1:3, and 2:2 stoichiometries, and there will be a broad distribution of cellular E_A_/E_D_ ratios between 0.3 and 3, especially if MCU-mCerulean and MCUB-mVenus are transfected at different ratios. Conversely, if MCU:MCUB composition is at a fixed stoichiometry, then the cellular E_A_/E_D_ ratios will reflect that stoichiometry as described for a single channel (0.33, 1, or 3).

**Figure 5.**
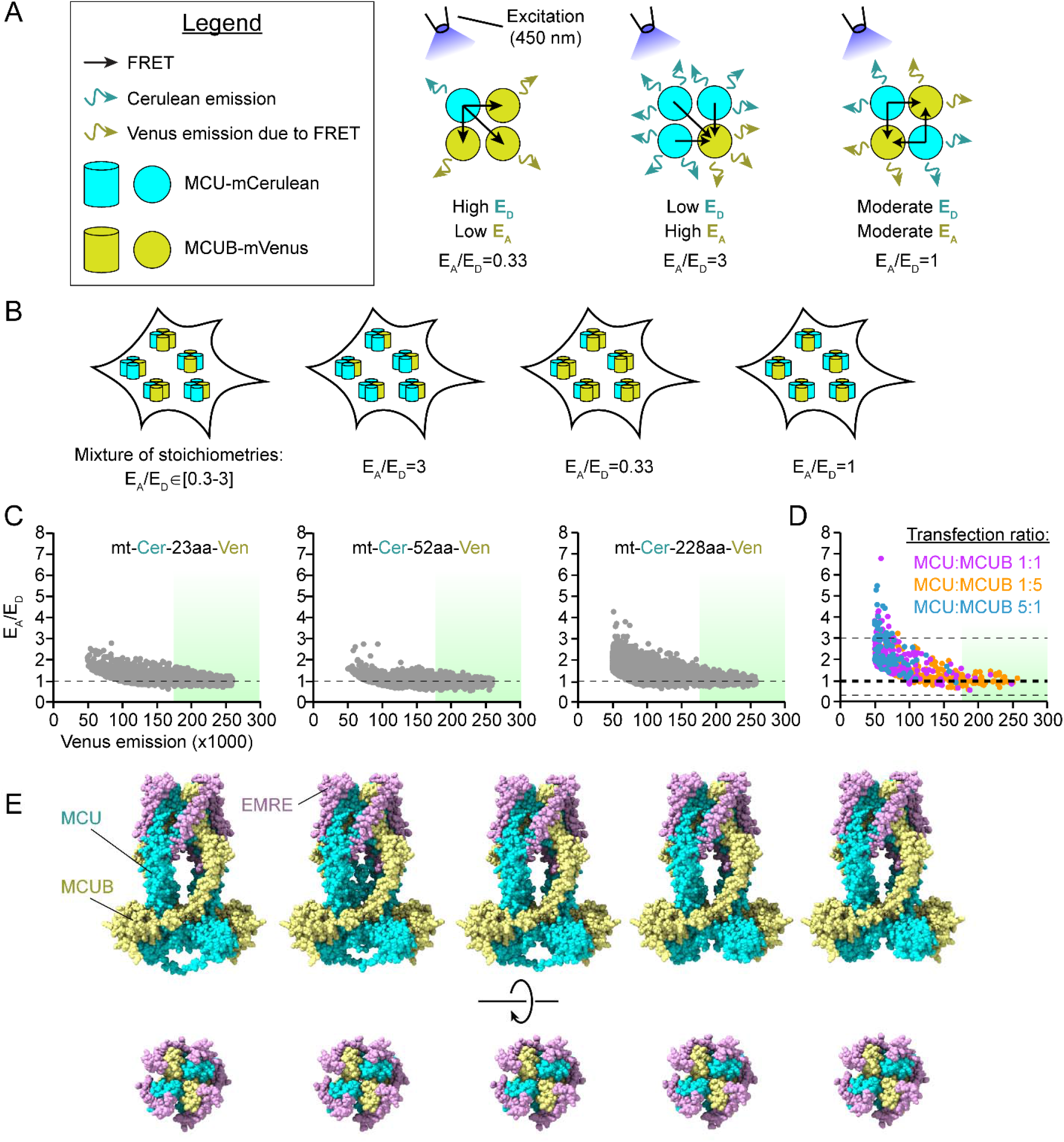
MCUB integrates into uniporter channels at a 2:2 ratio with MCU. **A**. Cartoons depicting how donor (E_D_) and acceptor (E_A_) FRET efficiency varies for uniporter channels containing different ratios of MCU:MCUB. MCU-mCerulean is the FRET donor, while MCUB-mVenus is the FRET acceptor. The E_A_/E_D_ ratio at high expression levels approximates the endogenous stoichiometry of each channel. **B**. Within each cell, uniporter channels can either have a variable stoichiometries (leftmost cartoon), leading to variable E_A_/E_D_ for each cell, or one of three fixed stoichiometries (right three cartoons), leading to fixed E_A_/E_D_. **C**. Proof of concept. Flow cytometry of transfected HEK-293T cells allows rapid collection of live cell FRET. Three different dimers of Cerulean-Venus with variable linker length, all show E_A_/E_D_ = 1 at high expression levels. The populations shown were selected from cells with high mCerulean expression. **C**. MCU/MCUB DKO cells were transfected with MCU-mCerulean and MCUB-mVenus at three different ratios, yet the distribution suggests a single E_A_/E_D_ ratio of 1, consistent with 2:2 MCU:MCUB. **D**. The top 5 AlphaFold3 predictions of 2:2:4 MCU:MCUB:EMRE all produce an abab architecture with alternating MCU and MCUB.

To implement a robust version of this protocol, we performed flow cytometry on MCU/MCUB DKO cells transfected with MCU-mCerulean and MCUB-mVenus at 1:5, 1:1:, and 5:1 DNA plasmid ratios. To validate the approach, we separately transfected HEK-293T cells with mitochondrially-targeted mCerulean and mVenus tandem dimers connected by linkers of variable length. For the mCerulean-mVenus dimers, although the FRET efficiency should vary inversely with linker length, with longer linkers producing less FRET due to greater distance between donor and acceptor, the 1:1 ratio of donor to acceptor should nevertheless produce an E_A_/E_D_ ratio of 1. For each cell, fluorescence from mCerulean, mVenus, and FRET was determined by correcting for cross-talk across excitation/emission channels, and these values were used to determine both E_D_ and E_A_, using formulas that estimate the FRET quenching of fluorescence for mCerulean (E_D_, E-FRET), and FRET sensitized emission from mVenus (E_A_, 3^3^-FRET) (Ben-Johny et al., 2016; S. R. Lee et al., 2016).

With this methodology, we found that the mitochondrial mCerulean-mVenus dimers showed E_A_/E_D_ ratios of ∼1 regardless of linker length, validating the FRET approach. Surprisingly, despite transfecting MCU and MCUB at different ratios, we found that cells also produced an E_A_/E_D_ value of ∼1, suggesting a 2:2 MCU to MCUB ratio. Taken together, these findings reveal that uniporter channels containing MCUB largely assemble at a fixed stoichiometry, and suggest a simple mechanism, expanded in the discussion, for MCUB-dependent down-titration of mitochondrial Ca^2+^ uptake.

With a 2:2 MCU to MCUB stoichiometry, the transmembrane portion of the channel can potentially assemble in two configurations. In one, two MCU subunits contact each other, with each one interacting with a single MCUB (e.g. MCUaabb). Conversely, the channel could assemble with each MCU subunit interacting with 2 MCUB subunits, and each MCUB subunit contacting 2 MCUs (e.g. MCUabab). FRET measurements cannot distinguish between these two scenarios, and structural data regarding the arrangement is lacking. To attempt to answer this question, we fed sequences for 2 MCU and 2 MCUBs (lacking the respective mitochondrial targeting sequences) into AlphaFold3, along with 4 EMRE subunits (Fig. 5E) (Abramson et al., 2024). Analysis of the permeation pathway for these structures was relatively unhelpful, as they assembled with similar shapes whether we used 4 MCU or 2 MCU:2 MCUB, and were narrower in all cases relative to published cryoEM structures (Wang et al., 2019). Nevertheless, the top five predicted channel architectures all assembled in an MCUabab configuration with high confidence, with each MCU subunit contacting two MCUB subunits and vice versa.

## DISCUSSION

In this study, we identify how the assembly and stoichiometry of MCUB allows dominant-negative regulation of the mitochondrial calcium uniporter. First, our findings revealed that MCUB does not express independently, but requires the presence of MCU for its expression. Second, the critical interactions allowing expression occur through the MCUB NTD, and replacement of this domain allows MCUB to express in isolation. Third, we observed that even multiple changes to MCUB transmembrane domains failed to restore Ca^2+^ uptake. Fourth, our data revealed that inhibition of mitochondrial Ca^2+^ uptake is proportional to the number of MCUB subunits incorporated into the pore. Finally, we identified the stoichiometry of MCUB-containing uniporters, showing that intact MCU and MCUB hetero-oligomerize in equal proportions (2:2 ratio) to form a tetrameric uniporter complex. Structural predictions suggest the architecture is such that each MCU subunit directly contacts two MCUB subunits, and vice versa (MCUabab conformation).

Our FRET assays revealed that MCU-MCUB containing uniporters were homogenously 2:2 within each cell. We cannot definitively discern with our data whether this distribution evolves by subunit exchange or during assembly. In the first case, uniporters with different MCUB subunit composition may exchange with each other and with MCU-only uniporters until the steady state distribution maximizing the number of channels containing the stable MCUabab architecture is achieved (Ernst et al., 2023). In the second case, if newly-made MCUB subunits integrate into channels faster than MCU, then it will only survive if there are MCU subunits nearby to bind, since MCUB-only channels do not express (Fig. 6A). We favor the second mechanism as more likely for several reasons: (1) once assembled into a channel, EMRE bridges interactions across neighboring MCU/MCUB subunits, which may limit the ability of these to exchange between different uniporters; (2) the MCUabbb concatemer we made expressed poorly, suggesting that channels with MCUB-MCUB interactions fail to survive; (3) given constant turnover of uniporter channels, it is unlikely steady-state would be reached under most circumstances, and we would expect cells to have more variable E_A_/E_D_ ratios, whereas our data is most consistent with a strict 2:2 ratio.

**Figure 6.**
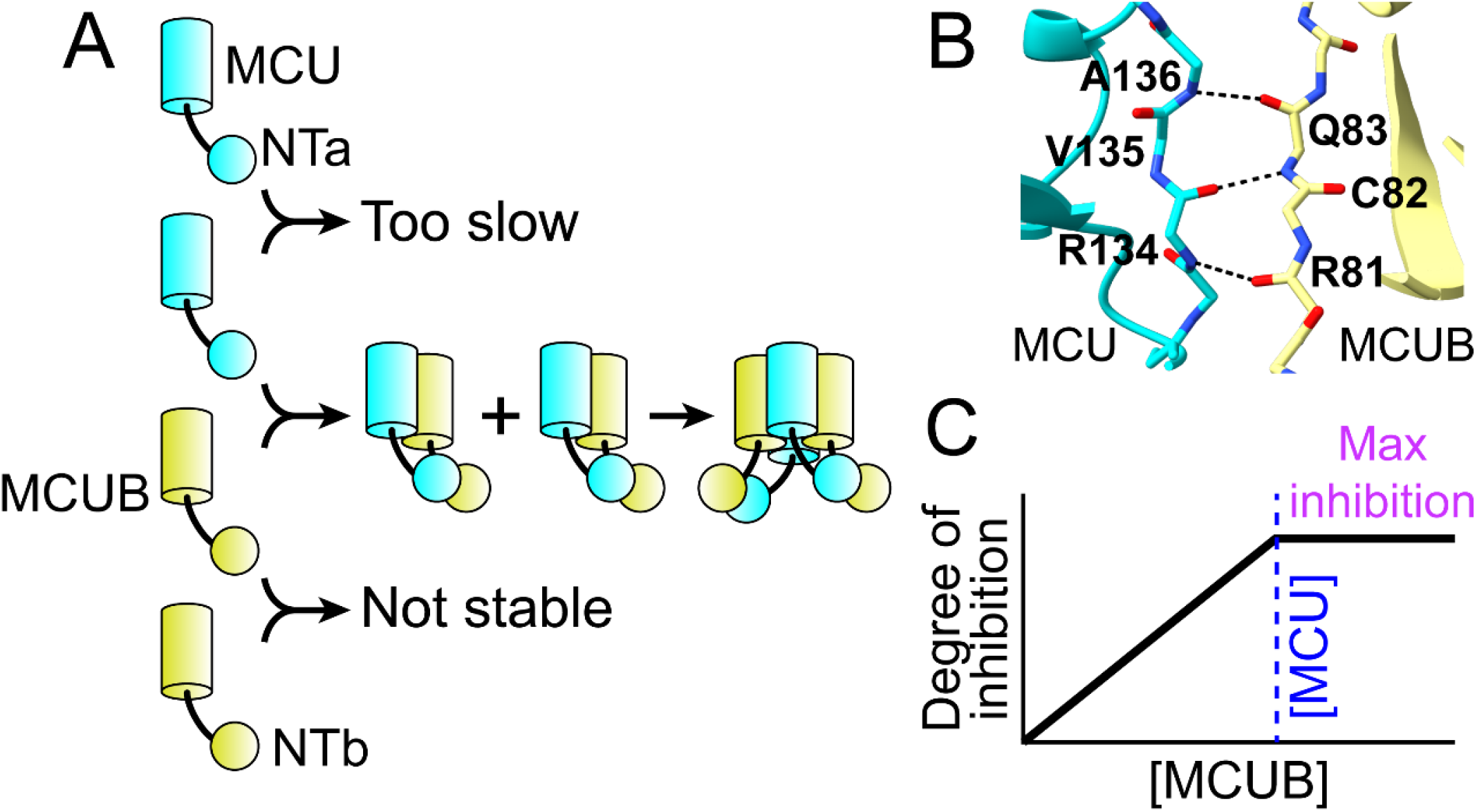
MCUB assembly into uniporter channels allows efficient and linear inhibition of Ca^2+^ uptake. **A**. Hypothetical MCUB assembly driven by NTa-NTb interactions. When MCUB is being produced, it preferentially integrates into uniporter channels driven by the stronger interaction between NTa-NTb compared to NTa-NTa. The preferential formation of oriented (asymmetric) dimers leads to an alternating abab architecture. NTb-NTb interactions fail to be stable, preventing the formation of MCUB-only channels. **B**. MCUB possesses the RCQ motif but lacks the RVA motif in its NTD (Fig. 2A), producing oriented assembly of MCU:MCUB dimers. The diagram is a close-up of NTa-NTb interactions in the highest confidence AlphaFold3-predicted structure. The NTa-NTb interaction is similar in the 4 other predicted structures. **C**. Because MCUB integrates into channels at a 2:2 ratio with MCU, cells can linearly tune Ca^2+^ uptake by controlling the ratio of nascent mitochondrial MCUB to MCU protein. When the MCUB concentration equals MCU (blue dashed line), maximal inhibition has been produced.

An elegant recent publication provides insight into how such a mechanism for preferential MCU-MCUB subunit incorporation into a channel may occur (Noble et al., 2024). The authors performed a detailed biochemical analysis of the purified, isolated MCUB NTb domain, and investigated how its stability was affected by the isolated MCU NTa protein. Two remarkable findings emerged. First, NTa-NTb dimers formed preferentially over NTa-NTa or NTb-NTb. The NTa-NTb heterodimer affinity was submicromolar and >60-130× stronger than either homodimer. Second, they also found that increasing NTb concentration led to instability in secondary structure and worsened solvent exposed hydrophobicity, suggesting that complexes containing only NTb tend to aggregate and are likely unstable. Such behavior was unusual and in contrast to NTa, which maintained stability at different concentrations.

To reconcile these findings with our data here, we propose the following mechanism for MCU-MCUB assembly. First, MCUB-MCUB dimers fail to form because of the instability of NTb-NTb driven interactions (seen also in the poor expression of MCUabbb concatemers, Fig. 4C), preventing the formation of MCUB-only and 1:3 MCU:MCUB channels. Second, when MCU monomers are available, these preferentially bind to available MCUB monomers due to the much greater stability of NTa-NTb interactions. This NTa-NTb dimer imposes an asymmetry on the structure. This occurs because, while NTa possesses both the RCQ and RVA motifs necessary for dimerization [see Fig. 3C in (Wang et al., 2019)], NTb possesses the RCQ but not the RVA motif (Fig. 2A, 6B). This asymmetry may thus orient two MCU-MCUB dimers so they can only assemble in an MCUabab pattern. Finally, the reason for the absence of 3:1 MCU:MCUB channels is somewhat unclear, since these concatemers express well and show Ca^2+^ uptake (Fig. 4C-E) but it may be that MCU subunit dimers fail to assemble stably when MCUB is present, since the NTa-NTb interaction is substantially preferred.

The fixed 2:2 stoichiometry combined with the inability of MCUB to express by itself, suggests a simple and efficient mechanism by which MCUB inhibits uniporter-mediated mitochondrial Ca^2+^ uptake. MCUB will preferentially titrate into uniporter channels, maximizing the number of complexes that are inhibited while not adopting inefficient configurations containing excess MCUB per channel (e.g. MCUB-only complexes, 1:3 MCU:MCUB). This predicts a linear relationship between the amount of mitochondrial MCUB protein and the degree of inhibition, with maximal inhibition occurring when [MCUB]=[MCU], allowing precise regulation of Ca^2+^ uptake (Fig. 6C).

Beyond stoichiometry, our data suggest MCUB integration reduces but does not abolish Ca^2+^ transport through the uniporter, addressing another unresolved question regarding its function. Raffaello et al. reported that addition of MCUB to MCU reduced the number of channels recorded without significantly altering electrophysiological properties, suggesting that adding MCUB fully abolished ion conduction. This was further supported by the finding that mutating two residues in MCU to their MCUB cognates (R252W and E257V) abolished Ca^2+^ uptake, perhaps by interfering with interactions with other uniporter subunits (Raffaello et al., 2013). However, an interesting feature of MCUB is the full preservation of the DIME sequence that coordinates Ca^2+^ in the pore. Why should this motif be preserved at all if the channel is non-conductive, given that point mutations in the motif are the most straightforward way to abolish Ca^2+^ transport while preserving complex formation? The preservation of this motif rather suggested that MCUB-containing channels may still support ion transport. This hypothesis was further supported by the finding by Raffaelo et. al. that MCUB reconstituted into planar lipid bilayers could transport Na^+^ in divalent-free conditions. Moreover, modeling studies by other groups revealed that a primary effect of MCUB incorporation into the uniporter may be to change the profile of the channel pore, implying that Ca^2+^ conduction may still be possible through the channel (Colussi & Stathopulos, 2023; Lambert et al., 2020).

To address this controversy, we created concatemers incorporating different ratios of MCU and MCUB, linked by a long flexible linker. The concatemers demonstrated a graded decrease in Ca^2+^ uptake that correlated with increased MCUB in each uniporter complex. Since all channels under these conditions would have one or more MCUB subunits present, these results reveal that MCUB incorporation reduces, rather than abolishes, Ca^2+^ uptake through the uniporter. Thus, allowing some Ca^2+^ transport to remain explains the preservation of the DIME motif in MCUB. To slow permeation without abolishing it is a harder optimization problem, and may explain why one or few mutations to MCUB were unable to restore Ca^2+^ flux. A possible caveat to these experiments was the finding that there was a low level of monomer still present in cells expressing the concatemers (Fig. 4C), and these may assemble to produce functional channels independent of the concatemers. However, this was primarily visible in the MCUaaaa and MCUabaa concatemers, whereas the preferred MCUabab and the more severe MCUabbb concatemers had little monomer evident, despite still showing some degree of Ca^2+^ uptake.

Our study has several limitations. All the work was performed in cell lines with heterologous expression, and differences in native systems may be noted. However, given the stability in uniporter behavior across tissues and cell lines, the mechanistic insight garnered here likely translates to the native setting. Nevertheless, we cannot exclude that in extreme conditions there may be some expression of MCUB-containing uniporter channels with other stoichiometries. In addition, our work with the MCU-MCUB concatemers suggests these can still conduct Ca^2+^. These assays relied on these concatemers adopting a normal configuration, and we cannot exclude the possibility that the long linkers may cause other rearrangements. However, it is unlikely that aberrant assemblies would conduct Ca^2+^, especially since even slight alterations to MCU lead to loss of Ca^2+^ uptake. Finally, our work focused on alterations to the transmembrane domains in inhibiting Ca^2+^ transport. Other mechanisms proposed, due to coupling between the N-terminal domains and the pore, or via altered interactions with accessory subunits, have not been tested here.

Following metabolic stressors such as ischemia-reperfusion injury, caloric restriction, or type 2 diabetes, MCUB expression increases 3-7 fold, with a concomitant decrease in mitochondrial Ca^2+^ uptake (Balderas et al., 2024; Cividini et al., 2021; Huo et al., 2020; Lambert et al., 2019). In the short term, such inhibition is protective against mitochondrial Ca^2+^ overload. However, chronic MCUB-dependent inhibition leads to a metabolic switch, with mitochondria becoming dependent on fatty acid oxidation over glycolysis, due to reduced activity or pyruvate dehydrogenase (Huo et al., 2023). Ultimately, such metabolic inflexibility may lead to impaired contractility and energetics. Our work here clarifies the mechanism for how transient changes in MCUB expression, driven by a range of these metabolic stressors, leads to direct, linear, and titratable reductions in mitochondrial Ca^2+^ uptake.

## Acknowledgments

We thank Dr. Vamsi Mootha for the kind gift of the MCU/MCUB knockout HEK-293T cell line. Support is from the National Institutes of Health (DP2ES032761 [YS], R01HL165797 [DC], R01HL141353 [DC]) and the Nora Eccles Treadwell Foundation (DC). The content is solely the responsibility of the authors and does not necessarily represent the official views of the National Institutes of Health.

## Data Availability Statement

The data that support the findings of this study are available from the corresponding author upon reasonable request.

